# Employing the intravesical delivery route to target on the kidney

**DOI:** 10.1101/2025.06.09.656495

**Authors:** Xinge Wang, Haiping Hu, Qiang Wang, Li Li, Xiao Z Shen

**Affiliations:** Department of Physiology, Zhejiang University School of Medicine, Hangzhou, Zhejiang, China; Department of Urology, Affiliated Zhejiang Hospital, Hangzhou, Zhejiang, China; Department of Laboratory Medicine, Affiliated Zhejiang Hospital, Hangzhou, Zhejiang, China; Department of Pharmacy, Affiliated Zhejiang Hospital, Hangzhou, Zhejiang, China

## Abstract

**Background and Objective:** There is an unmet demand for kidney-targeted non-invasive drug-delivery systems to enhance the therapeutic index and at the same time reduce extrarenal side effects. Here, we performed a proof-of-concept study to evaluate the validity and efficacy of intravesical delivery for kidney targeting in mice.

**Methods:** Probes with a wide range of molecular sizes were tested for their efficiency to reach the kidney via intravesical delivery in female mice. The therapeutic efficacy and side effects of pazopanib in treating a renal adenocarcinoma were compared between delivery via intravesical route vs. intraperitoneal route. Intravesical delivery of empagliflozin was tested for its action on the proximal tubules. Finally, intrapelvic infusion was examined as another retrograde route for targeting the kidney.

**Key findings and limitations:** Intravesical infusion was valid for a retrograde delivery of molecules up to 500 kDa to the kidney. Empagliflozin, an antagonist of sodium-glucose cotransporter 2 (SGLT2), could efficiently act on the proximal tubules via the intravesical route, and had an action on glucose excretion. In an orthotopic kidney carcinoma model, intravesical delivery of pazopanib was more efficacious in limiting tumor growth, accompanied with much milder adverse effects on extrarenal organs, compared to a systemic delivery when a same dose was administered. This was due to a higher intrarenal drug concentration achieved and a markedly lower level of drug leaked to the blood via intravesical delivery.

**Conclusions and Clinical Implications:** The intravesical delivery route has a great potential to benefit kidney therapeutics and research.

## INTRODUCTION

The prevalence of kidney diseases is increasing at an alarming rate globally.^1^ Therapeutics for kidney diseases are often limited by debilitating side effects. For example, patients with autoimmune kidney diseases are often prescribed with glucocorticoid which, however, is associated with systemic adverse reactions because of the nondiscriminatory distribution of the drug to organs/tissues outside of the kidneys.^2, 3^ The off-target side effects are particularly concerning chemotherapies, as the extrarenal toxic effects of chemotherapeutic agents have severely limited the dosages to be applied with.^4^ Therefore, there is an unmet demand for kidney-targeted non-invasive strategies for drug delivery to enhance the related therapeutic index relevant to kidneys and at the same time reduce the systemic side effects.

To address this concern, dramatic endeavors have been put for the development of nanotechnology- and macromolecular carrier technology-facilitated targeting systems to the kidneys, including liposomes, nanoparticles, antibody-modified carriers, and modifications enhancing endocytosis mediated by megalin and cubilin.^5-9^ However, these advancements require extensive chemical modification and still opt for a systemic route of delivery, e.g., via intravenous injection, which unavoidably affect extrarenal organs/tissues when accumulated by repeated application.

In a previous study, we demonstrated that intravesical administration of diphtheria toxin (DT) could efficiently deplete kidney macrophages in the *Cx3cr1*^CreERT2/+^:*iDTR* mice, sparing CX3CR1^+^ cells in other organs.^10^ This finding suggests that a retrograde route is practical and specific to deliver molecules from the bladder to the kidney. In the current study, we used mice to evaluate the kinetics and size-dependence of intravesical delivery for kidney-targeting. We show here that molecules even up to 500 kDa could reach proximal tubules via intravesical delivery. Moreover, we demonstrated that compared to a systemic route, the intravesical route was more efficient to deliver a chemotherapeutic agent for treating kidney tumor, at the same time inducing much milder adverse effects on other organs. Thus, the data of present study indicate a great potential of the retrograde urinary tract routes in kidney-specific delivery.

## METHODS

A detailed Methods section is available in the Supplemental materials.

## RESULTS

### Retrograde ascending of intravesically delivered probes to kidneys

To study the dynamics of retrograde ascending of molecules into the kidneys via the intravesical route, we first infused Rhodamine B, a saline-resolvable small fluorescent probe (479 Da), into the bladder of C57BL/6 female adult mice (8-12 weeks old). To do that, mice were first water-deprived for 4 hr to reduce urine production, followed by an infusion of 60 μl probe in saline to the bladder through a catheter (see Supplementary Methods). In 2 hr, fluorescence microscopy exhibited a significant number of probes had ascended into the kidney papilla (Figure 1A left), and the amount decreased afterwards (Figure 1B left). The probes also ascended into kidney medulla and cortex (including AQP1^+^ proximal tubules), although in a delayed manner relative to the papilla (Figure 1A middle and right). The peak of probe accumulation in the medulla and cortex appeared at 8 hr post intravesical infusion (Figure 1B middle and right), suggesting a continuous retrograde dispersion from the papilla to the upstream kidney during Hour 2-8 post infusion.

**Figure 1.**
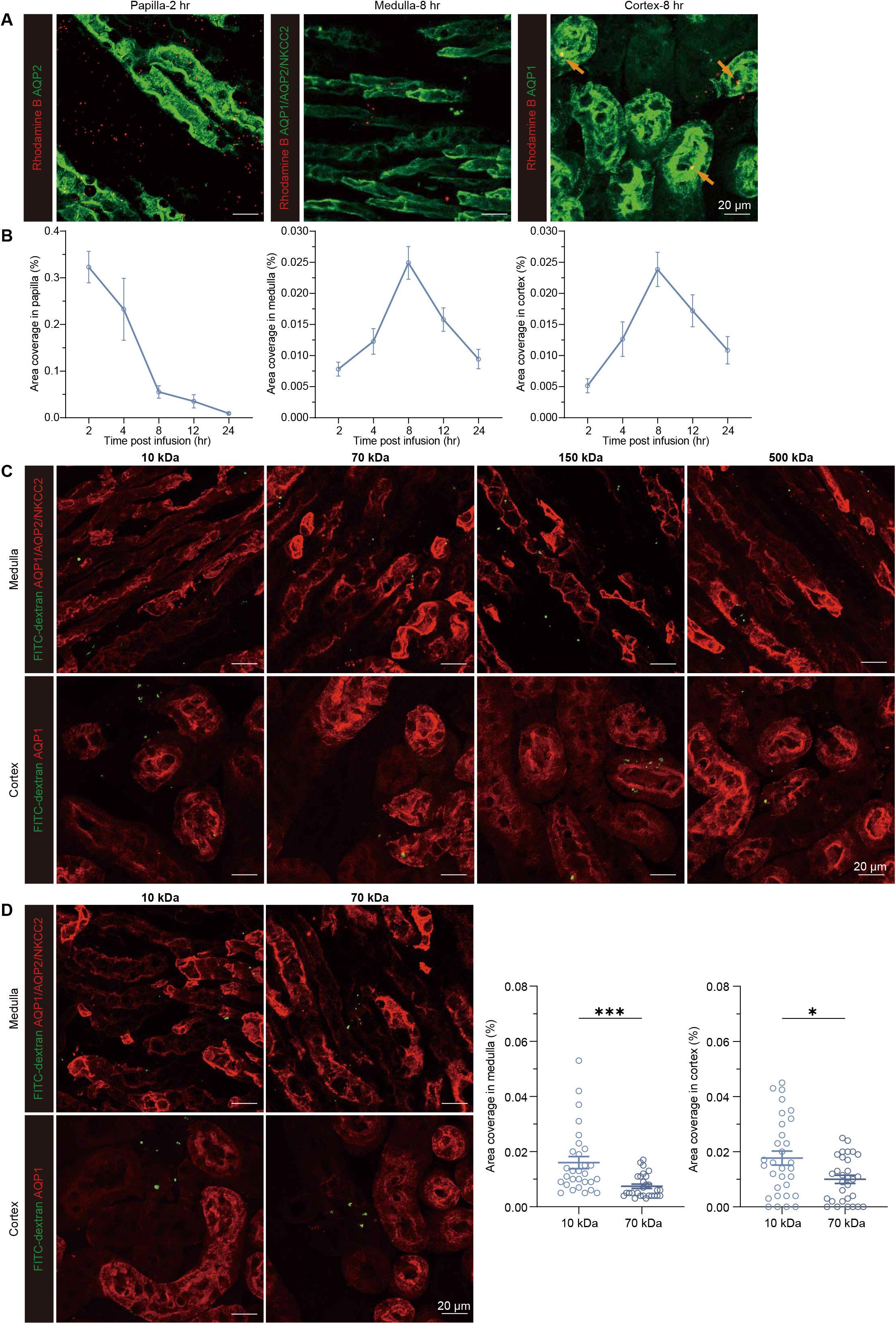
The dynamics of intravesical delivery of probes in different sizes to the kidney. (**A, B**) The representative images (A) and kinetics (B) of Rhodamine B in the renal papilla, medulla and cortex at the indicated time points after an intravesical infusion. Arrow, Rhodamine B in the proximal tubules. n = 6-9. (**C**) The representative images of 10, 70, 150, 500 kDa FITC-dextran in the medulla and cortex of kidney on hour 8 after an intravesical infusion. (**D**) The amounts of 10 and 70 kDa FITC-dextran in the medulla or cortex of the kidney were compared on Hour 8 after mice received an intravesical infusion of the same numbers of mole of 10 or 70 kDa FITC-dextran. Each dot indicates the quantification from one 319.5 × 319.5 μm^2^ fields of view (FOVs), n = 6. \**P*<0.05, \*\*\**P*<0.005 by two-tailed unpaired *t* test. Data are depicted as mean±SEM. Data are derived from at least 2 independent experiments.

We next investigated probes with larger sizes. It displayed that FITC-dextran with a molecular weight of 10, 70, 150, or even 500 kDa could reach both cortex and medulla of the kidney at hour 8 post intravesical administration (Figure 1C), demonstrating that the intravesical route was practical to deliver molecules with a wide range of size into the kidney. To interrogate whether size is a limiting factor for delivery efficiency to the kidney, we compared the ability of 10 and 70 kDa FITC-dextran ascending into the medulla and cortex. To this end, both of the 10 and 70 kDa FITC-dextran were prepared with an equivalent molar ratio of the fluorophore FITC and dextran. Equal molar amounts of these probes were individually infused intravesically and their relative quantities in the kidney cortex and medulla were compared after 8 hr (Figure 1D). It showed that smaller molecule appeared more efficient in retrograde ascending to the kidney. Substantiating this notion, intravesical infusion could barely deliver 0.5 μm fluorescent latex beads, a particle with size much larger than 500 kDa dextran, into the kidney (data not shown).

Of note, intravesical injection of 60 μl saline daily for 7 consecutive days did not inflict damages to the kidney, since no obvious apoptosis and inflammation occurred in the kidney, as manifested by staining of cleaved caspase-3 and expression of a panel of inflammatory cytokines, respectively (Figures S1A and S1B). Also, functional indices of kidney, including creatinine (CREA) and urea nitrogen (BUN) in blood, were normal after the intravesical regimen (Figure S1C).

### Intravesical delivery of chemotherapeutic agent is more therapeutically efficacious to treat renal adenocarcinoma with fewer adverse effects than systemic delivery

Next, we made comparison of the efficacy and side effects of a chemotherapeutic agent delivered via intravesical route vs. intraperitoneal route in a kidney tumor model. To this end, the renal adenocarcinoma (RENCA) cells were orthotopically implanted to female syngeneic BALB/c mice to create primary kidney tumors. The tumor cells were genetically labelled with luciferase (RENCA^luc2^), so bioluminescent imaging could be applied to measure the tumor size. Tumor sizes were first measured on Day 6 post implantation, and based on the tumor sizes, mice were divided into 3 groups to assure each group had a comparable average tumor burden. From Day 7, two groups were treated daily with pazopanib (437.5 Da, comparable to the size of Rhodamine B), a reagent previously shown to successfully suppress RENCA growth^11-13^, with one via intravesical route and another with intraperitoneal route. The third group were treated daily with vehicle as a control. After a 7-day treatment, i.e., on Day 14 post tumor implantation, tumor size was measured again (Figure 2A). It showed that while both groups with chemotherapeutics had smaller tumors compared to the control group, intravesical treatment could almost completely impede tumor growth, with a significantly lighter tumor burden at the end of treatment in comparison with intraperitoneal treatment (Figure 2B). Collaborating this finding, intrarenal concentration of pazopanib on Hour 8 post intravesical treatment (the time point when its peak concentration reached in the kidney, referring to the kinetics of Rhodamine B) was markedly higher (∼3 folds) than that in the mice treated via the intraperitoneal route (Figure 2C). Of note, there was no difference in intrarenal pazopanib concentration between Hour 2 and Hour 8 post intraperitoneal infusion, indicating that there was no overt gradual accumulation of pazopanib in the kidney after an intraperitoneal infusion.

**Figure 2.**
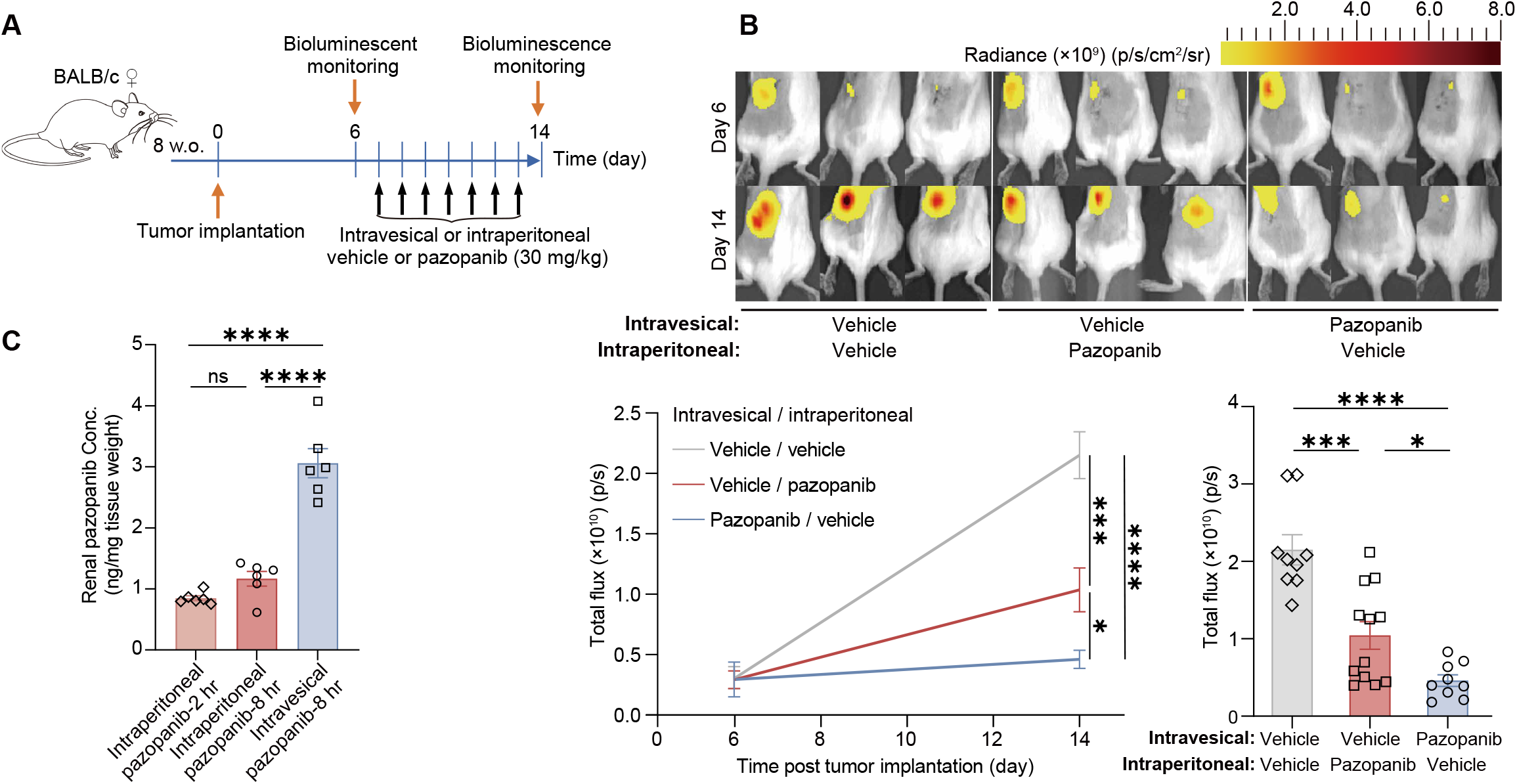
Therapeutic effects of pazopanib on treating renal adenocarcinoma via intravesical or intraperitoneal delivery. (**A**) Scheme illustrating the protocol of orthotopic implantation of RENCA tumor cells to BALB/c mice followed by a pazopanib treatment in a same dose but via different routes. (**B**) Tumor sizes with or without pazopanib treatment via intravesical or intraperitoneal route at day 6 and day 14 which represent the day before and after pazopanib treatment, respectively. (**C**) Mice received an intraperitoneal or intravesical infusion of pazopanib in a same dose. The concentrations of pazopanib in the kidneys were measured at the indicated time points. \**P*<0.05, \*\*\**P*<0.005, \*\*\*\**P*<0.001 by one-way ANOVA with Tukey’s multiple comparisons test. Data are depicted as mean±SEM. Data are derived from at least 3 independent experiments.

Therefore, we conclude that the peak concentration of drug obtained via the intravesical route is certainly higher than that obtained via the intraperitoneal route. Given the fact that peak concentration of chemotherapeutic agent at tumor loci is critical for its anti-tumor effect,^14^ our data suggest that the anti-tumor effect on kidney is superior with an intravesical delivery of chemotherapeutic agents over a systemic delivery of the same dose.

To evaluate the side effects of pazopanib via intravesical route vs. intraperitoneal route, we treated female BALB/c mice with pazopanib daily for 7 consecutive days (Figure 3A). Hypertension is a common side effect of pazopanib,^15-18^ and our previous work also showed a hypertensive effect of pazopanib in wild-type mice.^19^ We found here that one dose of pazopanib via intraperitoneal infusion could raise blood pressure and a 7-day intraperitoneal treatment further increased blood pressure (Figure 3B). In contrast, the mice treated with pazopanib via intravesical infusion did not develop hypertension even after 7 shots (Figure 3B). Blood cell counting showed that the mice with intraperitoneal treatment had reduced counts of white blood cells and platelets, which was not observed in the mice having intravesical treatment (Figure 3C the upper left and middle panels). Although both groups had no alteration in red blood cell count, hematocrit or hemoglobin concentration (Figure S2A), reticulocytes (circulatory immature red blood cells) decreased in number in the mice treated via intraperitoneal route but not in the counterparts treated via intravesical route (Figure 3C the upper right panel). These data altogether suggested that a systemic delivery but not an intravesical delivery of the chemotherapeutic agent would affect bone marrow hematopoiesis. Besides, a rise of biochemical markers indicative of malfunctioned liver (aspartate aminotransferase, AST, and alanine aminotransferase, ALT) and heart (lactate dehydrogenase, LDH) occurred in the mice treated via the intraperitoneal route but not in those treated via the intravesical route (Figure 3C the lower panels). Patients with pazopanib treatment barely reported side effects on kidney.^15, 16^ In alignment, blood CREA and BUN, did not increase in either of the pazopanib-treated groups (Figure S2B); also, there was no alteration of general architecture of the kidney in either of the groups (Figure S2C). To explain the reason for the much milder adverse effects via intravesical treatment vs. intraperitoneal treatment, we measured the concentration of pazopanib in blood plasma. Indeed, on Hour 8 post intravesical treatment when the highest level of intrarenal pazopanib was reached (referring to the kinetics of Rhodamine B), pazopanib concentration in blood plasma was significantly lower than that at Hour 8 post delivering via the intraperitoneal route (Figure 3D). Actually, the higher blood level appeared in the early time point, i.e., Hour 2, for the intraperitoneal delivery (Figure 3D), consistent with a previous report.^20^ As such, pazopanib leaked to the blood by the intravesical route was marginal (3.65%) relative to that reached via an intraperitoneal delivery. These data suggest that intravesical administration had minimal leakage to the blood and was much safer regarding systemic off-target adverse effects which usually limit the dosage applicable for many drugs.

**Figure 3.**
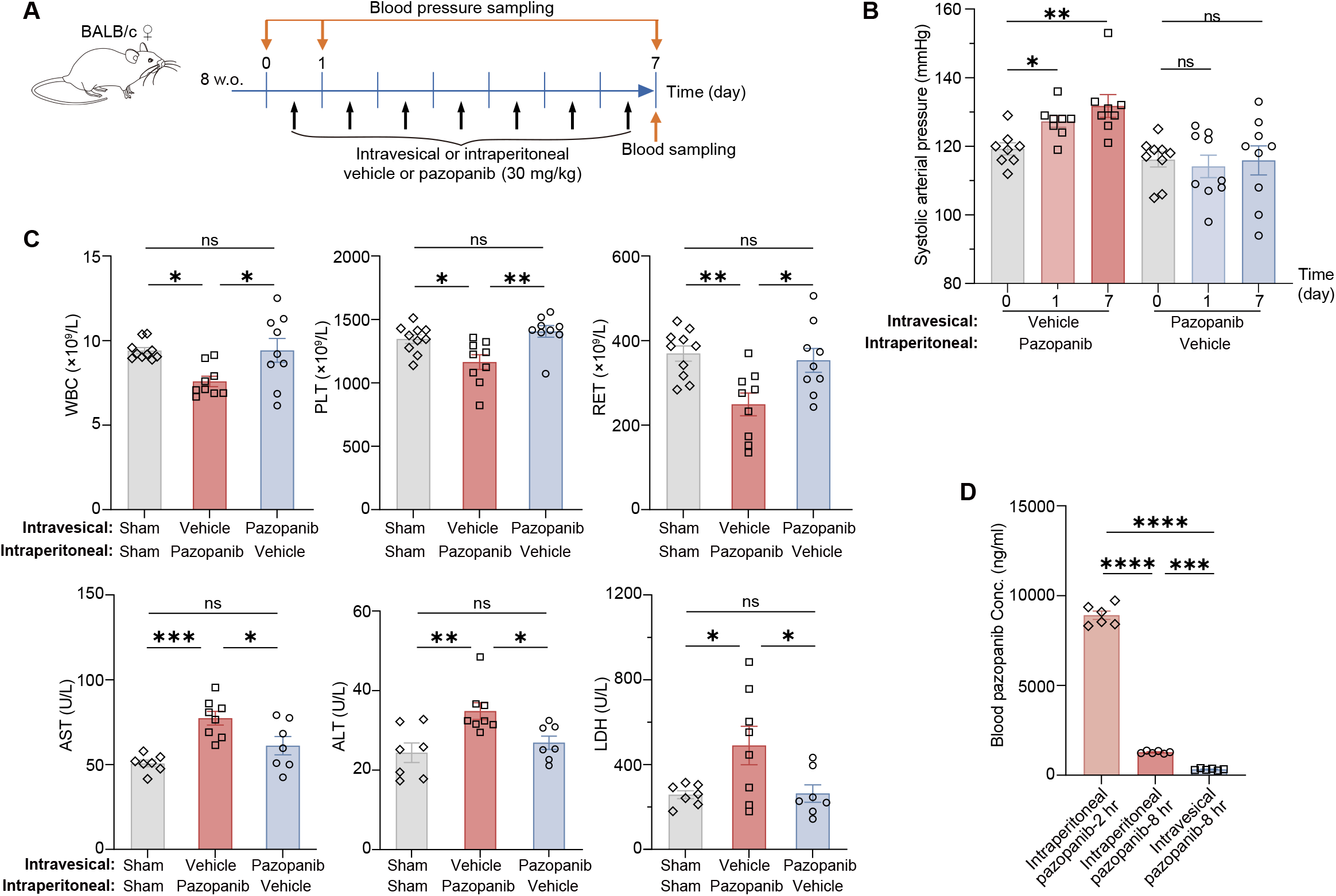
Side effects of pazopanib via intravesical or intraperitoneal delivery. (**A**) Scheme illustrating the experiments (B-D) to compare the adverse effects of pazopanib after delivered via intravesical route vs. intraperitoneal route. (**B**) Systolic blood pressure was measured at the indicated time points. (**C**) Counts of white blood cells, platelets and reticulocytes (RET) in the blood and blood levels of AST, ALT and LDH. (**D**) Mice received an intraperitoneal or intravesical infusion of pazopanib in a same dose. The concentrations of pazopanib in the blood were measured at the indicated time points. ns, not significant. \**P*<0.05, \*\**P*<0.01, \*\*\**P*<0.005, \*\*\*\**P*<0.001 by one-way ANOVA with Tukey’s multiple comparisons test. Data are depicted as mean±SEM. Data are derived from at least 3 independent experiments.

### Drug delivered via intravesical route can act on proximal tubules

Proximal tubules are the uppermost segment of the renal tubule system and are the major places where reabsorption of most molecules including glucose from the glomerular filtrate takes place. To examine whether drugs delivered via intravesical route can affect targets expressed on the proximal tubules, we tested empagliflozin, an antagonist of sodium-glucose cotransporter 2 (SGLT2) which mainly distributes in the proximal tubules and reabsorbs about 90% of the filtered glucose. Indeed, intravesical treatment with empagliflozin (451 Da) for a week could efficiently increase glucose excretion from the urine (Figure 4A left). Since SGLT2 is a sodium-glucose cotransporter, an accompanied over-excretion of Na^+^ was observed as well (Figure 4A middle). Also, given the increased glucose and Na^+^ in the urine which unavoidably elevated osmolality, more urine output was observed in the empagliflozin-treated mice in comparison to the control mice which had an intravesical treatment of vehicle (Figure 4A right). A previous study reported an acute (in one hour) effect of intragastrically administered empagliflozin on decreasing blood glucose level.^21^ To examine whether such an acute effect of empagliflozin could be achieved via the intravesical route, we performed blood glucose test on fasted mice. One intravesical shot of empagliflozin or vehicle was given to mice fasted for 6 hr and we monitored their glycaemia in the following 8 hr (Figure 4B). With a decreasing trend of blood glucose in all of the mice during the observation period (due to a continuous fasting), mice with the intravesical treatment of empagliflozin had a further downregulation of blood glucose level on Hour 6-8 post the treatment (Figure 4B). This timeline for empagliflozin to taking effect was consistent with our previous finding that Rhodamine B which has a similar molecular size to empagliflozin reached the highest concentration in the kidney cortex at around Hour 8 post intravesical infusion. Thus, these data demonstrate that therapeutic agents delivered via the intravesical route can take effects on the molecular targets arguably present in any segment of the renal tubule system.

**Figure 4.**
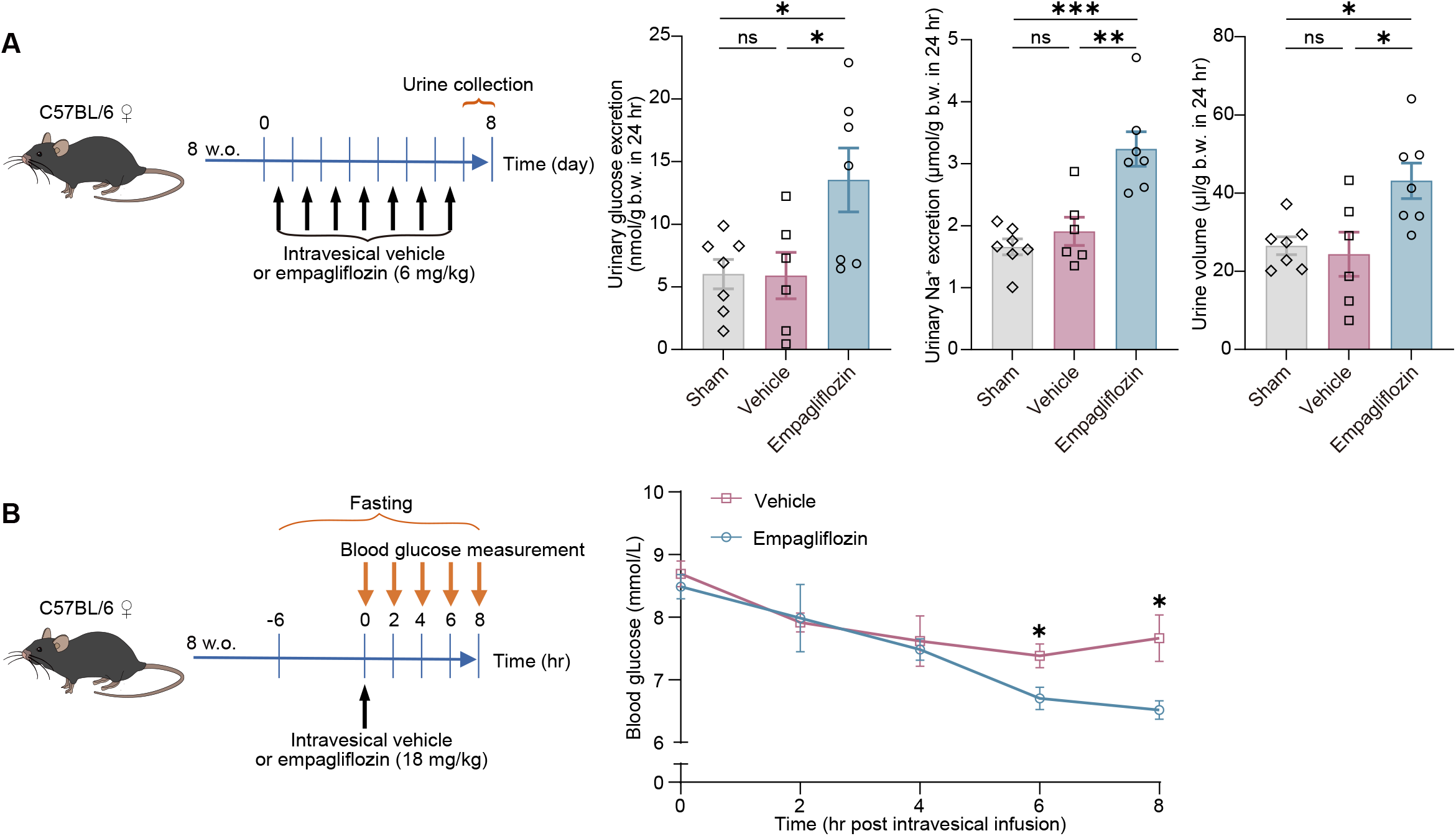
Intravesically delivered empagliflozin can block SGLT2 and affect glucose excretion. (**A**) The experimental protocol for an intravesical regimen of empagliflozin treatment (left). Urine outputs of glucose, Na^+^ and urine volume were measured after the regimen (right). (**B**) The experimental protocol for an acute intravesical treatment of empagliflozin after fasting (left). The blood glucose levels were measured at the indicated time points after the treatment (right). n = 7. ns, not significant. \**P*<0.05, \*\**P*<0.01, \*\*\**P*<0.005 by one-way ANOVA with Tukey’s multiple comparisons test in (A), two-tailed unpaired *t* test in (B). Data are depicted as mean±SEM. Data are derived from at least 2 independent experiments.

### Intrapelvic infusion is more efficient to deliver molecules into the kidney than intravesical infusion

Ureteral catheter has been widely used in clinic for multiple applications.^22, 23^ It can pass through the bladder and ureter and reach up to pelvis in human, suggesting that this device could be adapted for an application of non-invasive intrapelvic infusion. As such, intrapelvic infusion may be another useful route for kidney delivery in human. To interrogate the efficiency of intrapelvic delivery in targeting at the kidney, we made head-to-head comparison between intrapelvic and intravesical delivery of in delivering Rhodamine B in mice (Figure 5A). In this comparison, the dose for intrapelvic infusion was halved to that for intravesical infusion since the bladder connects two kidneys. Unfortunately, there is no ureteral catheter developed for mice. To do the intrapelvic injection, we surgically exposed the pelvis of one kidney and made an intrapelvic injection. Interestingly, we observed a similar kinetics of Rhodamine B reaching the medulla and cortex via intrapelvic infusion to intravesical infusion, as the peak appeared at around Hour 8 (Figure 5B). However, there were much more probes present in both the medulla and cortex in the intrapelvic group than the intravesical group at any given time point (Figure 5C). In addition, a large size (500 kDa) probe recapitulated the result that the intrapelvic route was more efficient over the intravesical route for a retrograde delivery to the kidney (Figure 5D).

**Figure 5.**
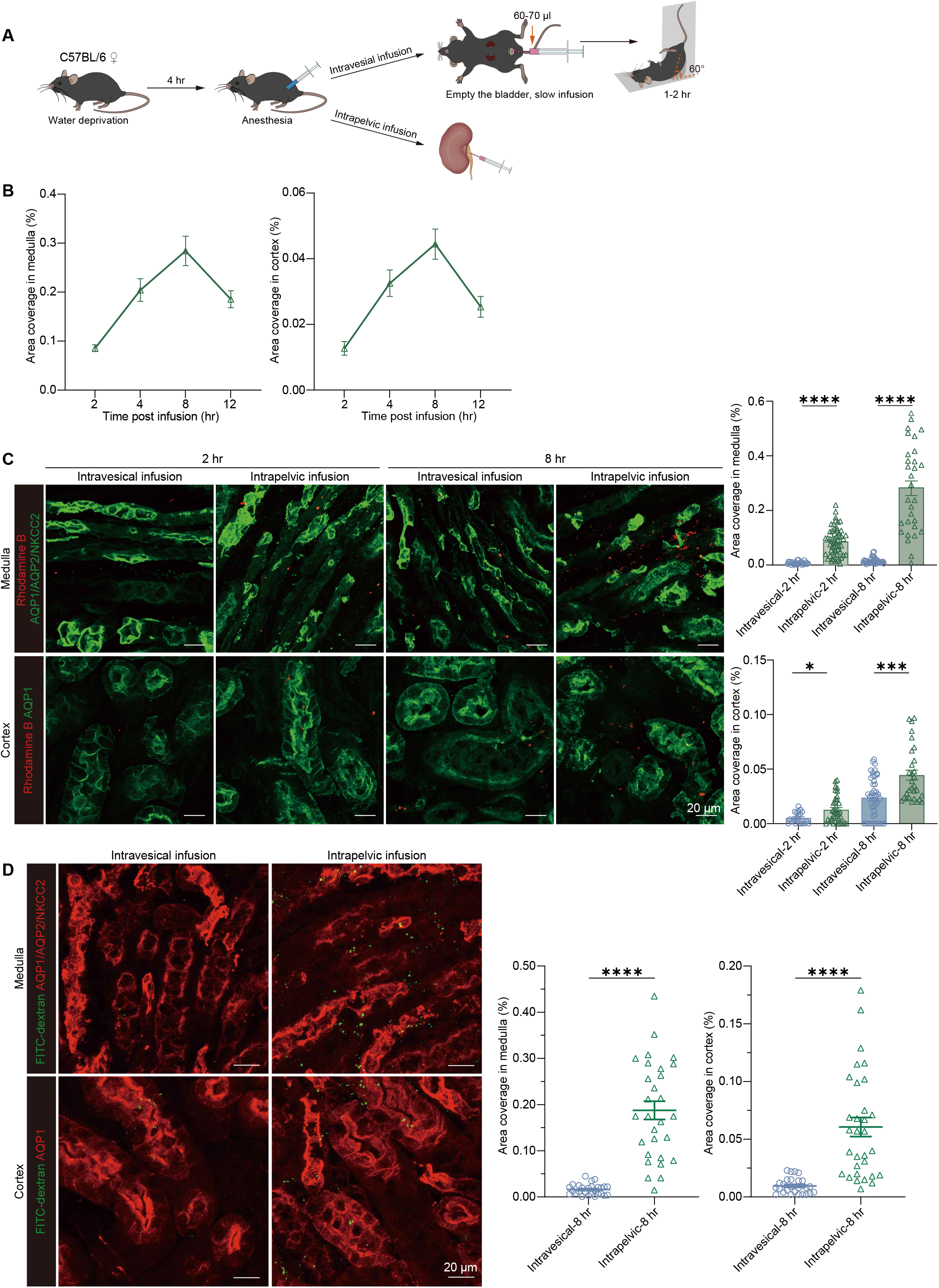
Intrapelvic delivery is another efficient route for targeting kidney. (**A**) Scheme illustrating the procedures for intravesical or intrapelvic infusion. (**B**) The dynamics of Rhodamine B reaching the medulla and cortex of kidney after an intrapelvic infusion. n = 6-7. (**C** and **D**) After an infusion of Rhodamine B (C) or 500 kDa FITC-dextran (D) via the intravesical or intrapelvic route, the abundance of probes in the cortex and medulla was measured at the indicated time points. Each dot indicates the quantification from one 319.5 × 319.5 μm^2^ FOVs. n = 6-9 for (C); n = 6 for (D). \**P*<0.05, \*\*\**P*<0.005, \*\*\*\**P*<0.001 by two-tailed unpaired *t* test. Data are depicted as mean±SEM. Data are derived from at least 3 independent experiments.

## DISCUSSION

Current treatment strategies for kidney diseases are generally based on systemic administration, which brings about unwanted toxicity and adverse effects on other organs/tissues. Indeed, the intolerance of side-effects is the major hurdle for a sufficient dosage and a continuous regimen.^4, 24-27^ In this context, retrograde urinary tract delivery was tested here as a non-invasive and kidney-targeting option. Intravesical delivery is usually used for treating bladder diseases, such as bladder cancers.^28, 29^ Moreover, ureter catheter is widely used to assist urine drainage from the upstream ureter and pelvis in conditions of urinary passage occlusion induced by various diseases including urolithiasis. Intravesical or intrapelvic delivery for targeting the kidney has not been seriously investigated before.

In the present study, we demonstrate that intravesical and intrapelvic infusion are able to deliver chemicals in a certain range of size to the upstream kidney. In particular, our preclinical results display a superior anti-tumor outcome with intravesical delivery of a chemotherapeutic agent relative to the same dosage applied via a systemic intraperitoneal delivery. Moreover, this was accompanied with much milder side-effects on extrarenal organs. These biochemical outcomes were supported by analysis of the concentrations of the chemotherapeutic agent in the blood and the kidney, as intravesical delivery resulted in a negligible level of the drug in the blood and a much higher level in the kidney relative to the systemic delivery. These data suggest that a better therapeutic outcome could be achieved via intravesical delivery of a higher dose via otherwise forbidden when given through a systemic delivery.

Benefited from the urinary tract delivery, drugs do not require specific modifications for kidney targeting. We show here that molecules with size up to 500 kDa could be delivered to the kidney via the urinary tract routes, suggesting the drugs which would be otherwise excluded by glomeruli filtration could reach and act on the tubule system by the urinary tract delivery. Thus, targeting the kidney by urinary tract delivery has great potential to expand drug usage and provide new angle to drug development for kidney diseases.

## AUTHOR CONTRIBUTIONS

Conceptualization, X.Z.S.; designing, X.W. and X.Z.S.; supervision, L.L. and X.Z.S.; imaging, X.W.; blood and urine biochemical analysis, Q.W.; all other experiments, X.W. and H.H.; mass spectrometry, L.L.; statistics, X.W.; validation, L.L.; writing, X.Z.S. All authors have read and agreed to the final version of the manuscript and are accountable for the entire study.

## DATA AVAILABILITY STATEMENT

All data are shown in the manuscript, and there is no restriction to provide any raw data.

## ACKNOWLEDGMENTS

We thank Jingyao Chen and Qiong Huang from the Core Facilities, Zhejiang University School of Medicine for the technical support. We also thank Dr. Yuancheng Weng for critical proofreading for this manuscript. This work was supported by National Natural Science Foundation of China (32170894 and 32470946 to X.Z.S) and Pioneer and “Leading Goose” R&D Program of Zhejiang(2025C02172 to L.L.).

