## Supplemental Figures 1-2 for "Employing the intravesical delivery route to target on the kidney"

#### Supplementary Material:

Supplementary Methods.

Supplemental figure 1. Intravesical infusion of saline does not make damages to the kidneys.

Supplemental figure 2. Effects of pazopanib administration on blood and kidney.

#### Supplementary Methods.

##### Mice

Female 8-12 w.o. C57BL/6 mice and BALB/c mice were purchased from Zhejiang Vital River Laboratory Animal Technology Co., Ltd. Mice were housed in specific-pathogen-free animal facility, with a 12-hour light/dark cycle. The experimental procedures were approved by Zhejiang University IACUC in accordance with the Guide for Care and Use of the Laboratory Animals. Authors have adhered to the ARRIVE guidelines for reporting animal research.

##### Reagents

The following probes were used in intravesical study: Rhodamine B (Solarbio, #R8040, 30 mg/kg), 10 kDa FITC-dextran (Sigma-Aldrich, #FD10S, 14.29 mg/kg), 70 kDa FITC-dextran (Sigma-Aldrich, #FD70S, 100 mg/kg), 150 kDa FITC-dextran (Sigma-Aldrich, #46946, 100 mg/kg), 500 kDa FITC-dextran (Sigma-Aldrich, #FD500S, 100 mg/kg). For intrapelvic study, the doses were halved accordingly.

For empagliflozin (MCE, #HY-15409), empagliflozin was first dissolved in DMSO (Coolaber, #CD4731C) and then diluted by 20 volumes in saline which contained 20% of sulfobutylether-β-cyclodextrin (Absin, #abs816067) by weight. its acute effect on glycaemia was tested by an intravesical delivery of 18 mg/kg, and its long-term effect on urine output was measured by daily intravesical delivery of 6 mg/kg for 7 consecutive days.

Pazopanib (Apexbio, #A3022) was first dissolved in DMSO and then diluted by 10 volumes in saline which contained 20% of sulfobutylether-β-cyclodextrin by weight. A dose of 30 mg/kg/day was used in this study.

**Intravesical injection**

Mice were first water-deprived for 4 hr to reduce urine accumulation. Afterwards, they were anesthetized with sodium pentobarbital (*i.p.*, 80 mg/kg) and kept on a homeothermic pad to maintain body temperature during the procedure. The bladder was gently manually massaged to be emptied, and then transurethrally inserted with a PE-10 catheter (ID: 0.28 mm, OD: 0.61 mm) through which 60 μl saline or indicated solution was slowly infused. After injection, the catheter with the syringe was retained for another 15 s to avoid a backflow. The mice were face-up lied with their posterior limbs elevated at a 60-degree till wakeup from the anesthesia (around 1-2 hr after the infusion).

**Intrapelvic injection**

Mice were anesthetized and kept on a homeothermic pad to maintain body temperature during the procedure. A unilateral left dorsal incision was made to expose left kidney. The adipose tissue surrounding the kidney was gently separated to expose the pelvis which locates in the renal hilus. An insulin syringe with a 30-gauge needle was used to inject 30 μl the indicated solution. The needle was kept in place for 10 s to forestall backflow and was then slowly pulled back. Gently press and swab the injection site to prevent any leakage. The kidney was gently returned to the abdominal cavity and the skin was sutured.

**Immunohistochemistry**

Mice were deeply anesthetized by 3% isoflurane and transcardially perfused consecutively with cold PBS followed by 4% paraformaldehyde (PFA). Kidneys were then isolated, fixed overnight in 4% PFA, and dehydrated in PBS containing 30% sucrose at 4 °C for 24 hr. Frozen sections (30 μm thick) were made on a cryostat (Leica, SM2010R) at -20 °C. Sections were blocked for 2 hr at room temperature with the blocking buffer (5% donkey serum in PBS). The sections were then incubated with a primary antibody in an antibody dilution buffer (0.25% Triton X-100 and 1% BSA in PBS) overnight at 4 °C, followed by incubation with secondary donkey antibodies in the antibody dilution buffer for 2 hr at room temperature in dark. After staining, sections were mounted on slides with DAPI Fluoromount-G anti-fade medium (Southern Biotech, #0100-20).

The following antibodies were used: rabbit anti-NCC (Millipore, #AB3553, 1:250); rabbit anti-SLC12A/NKCC2 (Abcam, #ab191315, 1:500); rabbit anti-AQP1 (Millipore, #AB2219, 1:2,000); rabbit anti-AQP2 (Abcam, #ab199975, 1:1,000); donkey anti-rabbit IgG AF488 (Abcam, #ab150073, 1:1,000); donkey anti-rabbit IgG AF568 (Abcam, #ab175470, 1:1,000).

**Confocal imaging**

After mounting on glass slides, confocal Z stacks were captured using the LSM 900 confocal microscope (Carl Zeiss, Germany) equipped with 20×/0.8 NA objective lens and processed with Zeiss ZEN (Blue edition) software. For fractional area quantification analysis, 26-μm-thick Z stacks images (1024 × 1024 pixels resolution) were recorded. Images were displayed as maximum-intensity projections of 25-μm-thick Z stacks recorded in 25 sections. Images were analyzed by ImageJ software (v1.54g, NIH, USA). The fractional areas were defined as the fraction of fluorescence positive areas. In brief, the sections from papilla, medulla or cortex, were analyzed based on the ratio of the calculated area of fluorescence-positive to the entire images (319.45 × 319.45 $\mu m^2$ area), and 5-6 fields from 2 different sections of each mouse were analyzed.

**Cell culture**

Murine renal adenocarcinoma cell line $RENCA^{luc2}$ (MeisenCTCC, #CTCC-0513-Luc2) was cultured in RPMI medium 1640 (Gibco, #C11875500BT) with 10% FBS, 1% HEPES and 1% penicillin-streptomycin in a humidified incubator at 37 °C with 5% $CO_2$.

**Orthotopic Renca-luc2 cell carcinoma model**

The mice were subjected to anesthesia and positioned in the right lateral decubitus posture. The surgery area was depilated and disinfected. A 1-cm longitudinal incision was made directly above the left kidney. The kidney was then squeezed out through the incision. $5\times10^5$ $RENCA^{luc2}$ cells suspended in 25 µl of PBS on ice were implanted orthotopically into the renal cortex of female BALB/c mice by an insulin syringe fitted with a 30-gauge needle. Afterwards, the syringe was kept in place for 10 s to forestall backflow and slowly pulled back. The muscle and skin were then sutured, followed by disinfection. The mice were kept warm until they recovered consciousness.

***In vivo* bioluminescent imaging**

To monitor and quantify tumor development, luciferase signals were detected on day 6 and 14 post $RENCA^{luc2}$ implantation. Mice were anaesthetized with 3% isoflurane and *i.p.* administered with 100 µl luciferin (MeisenCTCC, #CTCC-LUC-002) 9 min prior to imaging. Mice were placed with its dorsal side up. Imaging was performed using the IVIS Spectrum Imaging system (PerkinElmer, USA; 0.5 s, f/1, bin: 8, field of view: 23.4 cm) and bioluminescent signals were recorded as radiance (photons/s/$cm^2$/steradian). The tumor burden was measured by total flux (photons/s) within a defined region of interest (ROI) above left kidney using living image software (v4.3.1, PerkinElmer, USA).

**Measurement of arterial blood pressure by tail-cuff recording**

Systolic arterial pressure was determined in conscious mice using a computerized non-invasive tail cuff system (BP-2000 Series II, Visitech Systems, USA). Each mouse was trained for at least 5 consecutive days prior to data acquisition. Blood pressure was measured 25 times per session. Systolic arterial pressure was determined by averaging 20 measurements, with the first 5 readings discarded to minimize variability. The tracing waves were manually reviewed to verify proper blood pressure determination.

**Metabolic studies**

Urine was collected in a metabolic cage for 24 hr. Samples were then centrifuged at 12,000 g at 4℃ for 20 min and the supernatants were collected. Urine concentrations of sodium and glucose were measured by a chemistry analyzer Cobas c702 (Roche, Switzerland).

**Acute glucose-lowering test**

C57BL/6 female mice were fasted for 6 hr with free access to water. They were then intravesically administered with empagliflozin. Blood samples were obtained from the tail vein via carefully cutting off 1-2 mm of the tail tip with a sharp blade at 0, 2, 4, 6, 8 hr and the glucose levels were measured using a glucometer (Yuwell, #590, China). Always wipe off the first drop of blood to avoid hemolysis or contamination with interstitial fluid before taking new blood samples for blood glucose test. Food was withheld throughout the study.

**Plasma biochemistry and hematological analysis**

Mice were anaesthetized by inhaling 3% isoflurane. Blood was collected via retro-orbital bleeding using a capillary tube pre-treated with EDTA. Plasma was prepared by centrifugation at 1,500 g for 15 min at 4 °C, and the supernatant was collected for analyses by chemistry analyzer Cobas c702 (Roche, Switzerland). For quantification of blood cells, whole blood samples were analyzed by fully automated 5-part differential haematology analyser XN-1000V (Sysmex, Germany).

**Liquid chromatographic mass spectrometry (LC-MS)**

Before LC-MS analysis, blood was collected via cardiac puncture before perfusion, and kidney tissues were harvested after perfusion with ice-cold PBS. For the plasma samples, the plasma was separated by centrifuging the blood with EDTA at 1,500 g for 20 min at 4 °C. For the kidney samples to be tested, 1 ml acetonitrile and 1 ml deionized water were added to each gram of tissue. Kidney tissues were then homogenized at -10 °C using a high-speed low- temperature tissue homogenizer (Servicebio, #KZ-III-F, China) equipped with 3 cycles of 45 s processing at 60 Hz and each cycle followed by a 15 s cooling interval. Samples were then centrifuged at 12,000 g at 4 °C for 20 min and the supernatant was collected.

The supernatant was vacuum dried and reconstituted in 200 μl mobile phase consisting of (A): water (0.1% formic acid) and (B): acetonitrile, (88% B). 10 μl of solution was used each time for LC-MS analysis. Quantitative analysis was performed with Agilent 1260 HPLC chromatographic system (Agilent Technologies, USA) coupled to API4000 mass spectrometer (AB Sciex, USA) operated in positive electrospray ionization (ESI) through multiple reaction monitoring (MRM) mode. Data were analyzed with Analyst 1.6.3 software Hotfixes® (AB Sciex, Ontario, Canada). Kidney samples were normalized according to the dissected tissue weight.

**Statistics**

Statistical analysis was performed with Prism 8.0 (GraphPad). Data were shown as mean±SEM. One-way ANOVA with Tukey's multiple comparisons test was used to compare multiple groups. Two-tailed unpaired *t* test was used to compare two groups. P values of $<0.05$ were considered significant.

### Supplementary Figure 1

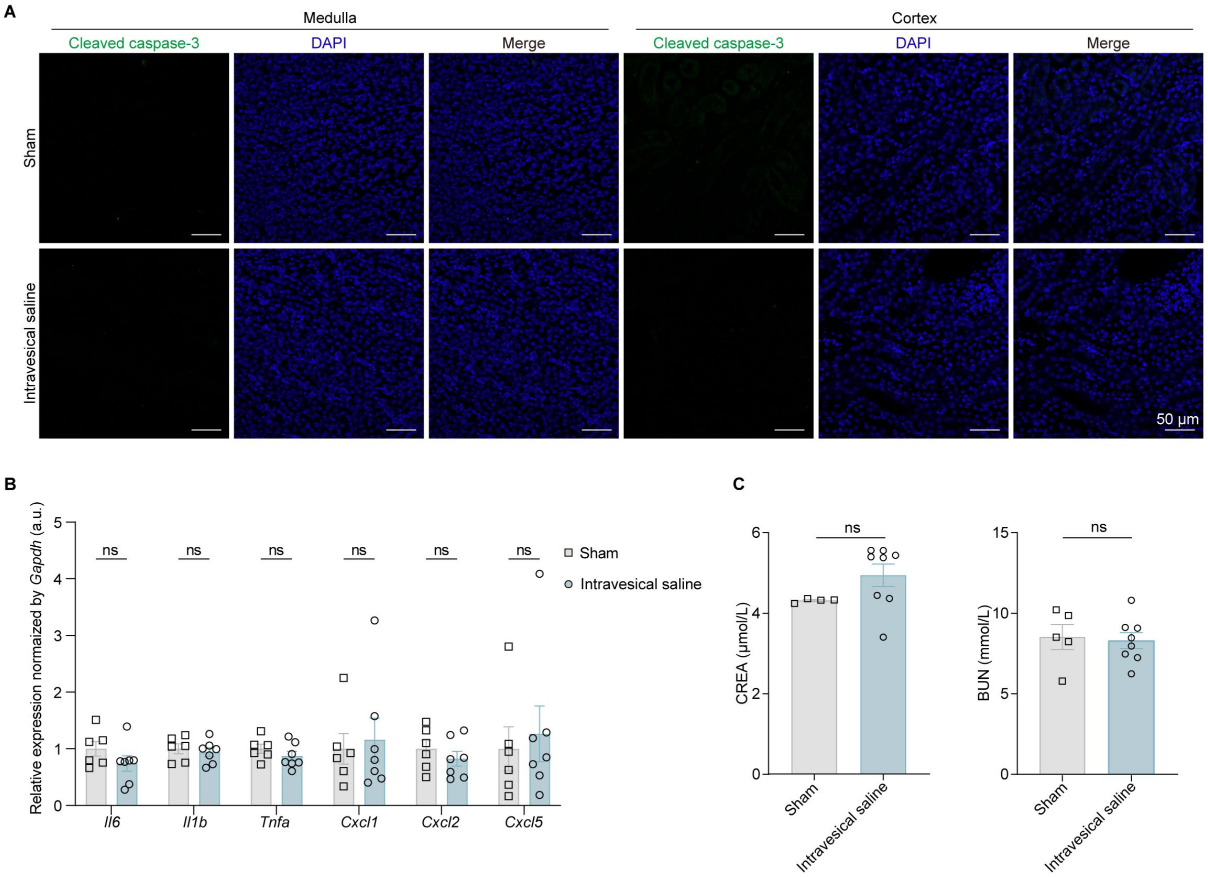


**Supplemental figure 1. Intravesical infusion of saline does not make damages to the kidneys.** Mice received daily infusion of 60 μl saline via the intravesical route for 7 consecutive days. (A) Immunohistostaining of kidney for cleaved caspase-3 was examined. (B) The proinflammatory cytokine expression in the whole kidneys were examined by RT-PCR. (C) Blood creatine (CREA) and urea nitrogen (BUN) concentrations were examined. ns, not significant by two-tailed unpaired t test. Data are depicted as mean±SEM. Data are derived from at least 2 independent experiments.

### Supplemental Figure 2

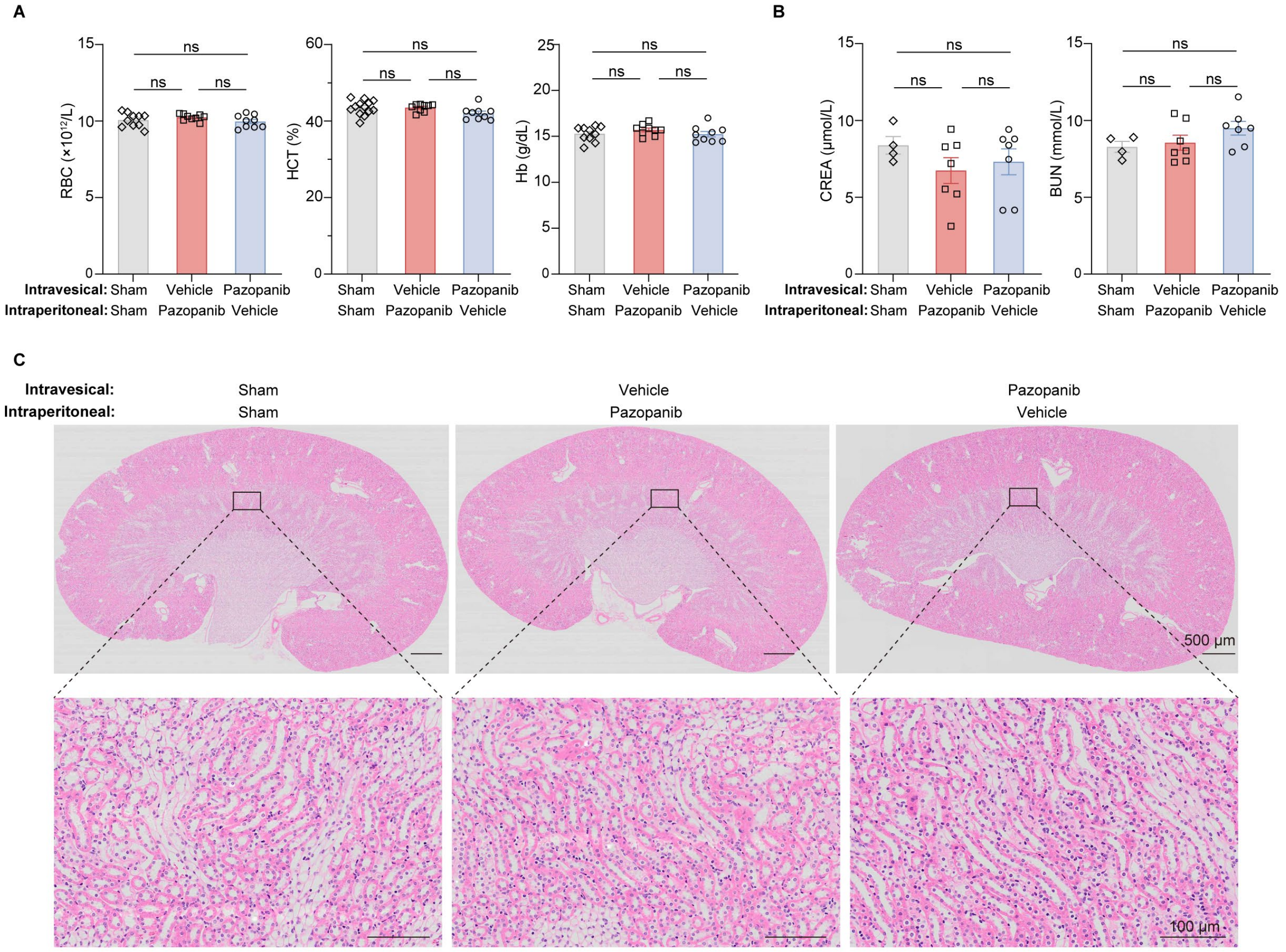


**Supplemental figure 2. Effects of pazopanib administration on blood and kidney.** Naive BALB/c mice received intraperitoneal or intravesical infusion of a same dose of pazopanib or were sham operated according to the protocol shown in Figure 2D. (A) Counts of red blood cells (RBC), hematocrit (HCT) and hemoglobin (Hb) concentration, and (B) creatine (CREA) and urea nitrogen (BUN) concentrations in the blood were measured. (C) A general histological structure of the kidney was evaluated by H&E staining. ns, not significant by one-way ANOVA with Tukey's multiple comparisons test. Data are depicted as mean±SEM. Data are derived from at least 3 independent experiments.
